# Variance misperception under skewed empirical noise statistics explains overconfidence in the visual periphery

**DOI:** 10.1101/2021.01.08.425966

**Authors:** Charles J. Winter, Megan A. K. Peters

## Abstract

Perceptual confidence typically corresponds to accuracy. However, observers can be overconfident relative to accuracy, termed ‘subjective inflation’. Inflation is stronger in the visual periphery relative to central vision, especially under conditions of peripheral inattention. Previous literature suggests inflation stems from errors in estimating noise, i.e. ‘variance misperception’. However, despite previous Bayesian hypotheses about metacognitive noise estimation, no work has systematically explored how noise estimation may critically depend on empirical noise statistics which may differ across the visual field, with central noise distributed symmetrically but peripheral noise positively skewed. Here we examined central and peripheral vision predictions from five Bayesian-inspired noise-estimation algorithms under varying usage of noise priors, including effects of attention. Models that failed to optimally estimate noise exhibited peripheral inflation, but only models that explicitly used peripheral noise priors -- but used them incorrectly -- showed increasing peripheral inflation under increasing peripheral inattention. Further, only one model successfully captured previous empirical results which showed a selective increase in confidence in incorrect responses under performance reductions due to inattention accompanied by no change in confidence in correct responses; this was the model that implemented Bayesian estimation of peripheral noise, but using an (incorrect) symmetric rather than the correct positively skewed peripheral noise prior. Our findings explain peripheral inflation, especially under inattention, and suggest future experiments that might reveal the noise expectations used by the visual metacognitive system.

**Significance:** Perceptual confidence can dissociate from accuracy in peripheral visual perception, a phenomenon known as peripheral inflation. No previous model has considered how this phenomenon may arise from metacognitive noise estimation which depends on empirical noise statistics. Here we simulate Bayesian-inspired noise estimation algorithms to show that the system’s erroneous beliefs about distributions of noise in the visual periphery can explain the occurrence of peripheral inflation, including how inflation varies with attentional manipulations in surprising ways. Our results explain why peripheral inflation occurs by positing a new Bayesian metacognitive noise estimation mechanism, paving the way for future psychophysical studies.

## A. INTRODUCTION

Great progress has been made towards understanding the internal statistical models that guide our perceptual decision-making and corresponding confidence ratings. When we make perceptual decisions about the world around us, they are accompanied by a metacognitive sense of certainty, or confidence, in whether those percepts are correct. Confidence ought to depend on the strength (signal) and reliability (noise) of the evidence to make the decision, i.e., should correlate with these measures (Fleming & Daw, 2017; Green & Swets, 1966; Macmillan & Creelman, 2005; Pouget et al., 2016); however, sometimes it does not (Fetsch et al., 2014; Koizumi et al., 2015; Maniscalco et al., 2016, 2020, 2021; Morales et al., 2020; Peters, Fesi, et al., 2017; Rahnev, Bahdo, et al., 2012; Rahnev et al., 2015; Rahnev, Maniscalco, et al., 2012; Rounis et al., 2010; Samaha et al., 2016, 2017; Zylberberg et al., 2012, 2016) or does so but not in a Bayesian optimal way (Adler & Ma, 2018; Denison et al., 2018); these cases suggest a need for innovation in models of perceptual confidence. Here, we sought to determine how natural scene statistics about noise distributions in the visual field might explain why confidence does not always perfectly track performance -- beyond a supposition that metacognition might simply involve “more noise” over and above noise in first-order (Type 1) estimates (Maniscalco & Lau, 2016).

A particularly intriguing case in which confidence and accuracy can dissociate is the phenomenon of subjective inflation, which is particularly evident in the visual periphery: subjective feelings of decisional confidence or visibility for unattended stimuli in the visual periphery are higher than those for stimuli in the center of the visual field for both complex and simple stimuli when task accuracy is matched (Knotts et al., 2020; Li et al., 2018; Odegaard et al., 2018; Rahnev et al., 2011; Solovey et al., 2014). In other words, observers appear ‘overconfident’ in their peripheral percepts relative to central percepts of the same signal reliability (Ehinger et al., 2017; Gloriani & Schütz, 2019; Hess et al., 2008; Rosenholtz, 2016). One explanation given for these findings is that observers’ Type 2 (metacognitive) criteria are fixed, such that under conditions of increased noise they do not adapt to the changing noise statistics of the environment, leading to a higher proportion of “high confidence” responses under a signal detection theoretic framework (Li et al., 2018; Rahnev et al., 2011; Solovey et al., 2014). However, the degree to which such Type 2 criteria are fixed remains open to some debate, as others have suggested that confidence judgments *are* sensitive to changing environmental or attentional conditions (Adler & Ma, 2018; Denison et al., 2018), albeit not in a Bayes-optimal manner. Importantly, it also remains unexplored why such explanations would provide stronger subjective inflation due to attentional manipulations in the visual periphery over central vision.

Other series of studies have suggested a conceptually analogous explanation: that illusions of confidence can be bidirectional, with both over- and under-confidence exhibited by the same system, due to a similar “fixed” estimate of noise, i.e. a “misperception” of variance (Gorea & Sagi, 2001; Peters, Fesi, et al., 2017; Zylberberg et al., 2014). That is, one element of the metacognitive system’s capacity to judge decisional accuracy is its ability to judge noise, i.e. signal reliability; by knowing a signal’s or representation’s noisiness, the observer can then set confidence criteria or engage in Bayesian inference appropriately. In cases of variance misperception, rather than directly measuring the reliability of its internal signals and using such an optimal measure in metacognitive calculations, the observer instead uses a heuristic -- a fixed, ‘typical’ level of noise applied unilaterally to metacognitive evaluations in the current task regardless of actually varying noise conditions (Gorea & Sagi, 2001; Peters, Fesi, et al., 2017; Zylberberg et al., 2014) or otherwise possesses mistaken beliefs about noise levels ((Drugowitsch et al., 2014; Fleming & Daw, 2017); see also (R. A. Adams et al., 2013)). It has been shown that such variance misperceptions can explain bidirectional confidence errors relative to decisional accuracy, and are related to the fixed metacognitive (Type 2) criteria also mentioned above (Li et al., 2018; Rahnev et al., 2011; Solovey et al., 2014).

Despite these potential answers due to fixed confidence criteria or variance misperception, a core explanation for why peripheral inflation occurs remains elusive for at least two reasons. First, it remains unclear how such heuristic ‘typical noise level’ judgments might be achieved: by what mechanism does the system “pick” a typical level of noise, and how is it used to set confidence criteria? And second, as with fixed criteria, variance misperception alone doesn’t necessarily explain why illusions of confidence appear to be easier to achieve in the periphery relative to central vision: on average, variance misperception (i.e., errors in noise beliefs) should lead to similar over- or under-confidence in general for both central and peripheral vision, unless the system were differentially applying misperceived variance in peripheral versus central vision. Instead, we see two main trends: subjective inflation in the periphery happens more than in the center under normal visual conditions (Knotts et al., 2020; Li et al., 2018; Odegaard et al., 2018; Rahnev et al., 2011; Solovey et al., 2014) (i.e., not in the dark; (Gloriani & Schütz, 2019)), and on average there is no inflation or deflation in central vision (Zylberberg et al., 2014).

Here, we sought to develop a simple explanation for these phenomena by combining these theoretical explanations with hypotheses about their potential source in empirical distributions of internal noise, in the same vein as the well-known influence of natural scene statistics on visual perception in general. Generally, the visual system and its perceptual computations are sensitive to external natural scene statistics (W. J. Adams et al., 2004; Girshick et al., 2011; Peters et al., 2015; Stocker & Simoncelli, 2006; Weiss et al., 2002)(W. J. Adams et al., 2004; Girshick et al., 2011; Peters et al., 2015; Stocker & Simoncelli, 2006) and human observers adapt their internal models to such external environmental statistical properties (Seriès & Seitz, 2013) -- making use of knowledge that cardinal orientations are more common, light typically comes from above, or motion tends to be slow and smooth, for example. Knowledge of these external natural scene statistics drives the formation of prior beliefs in a Bayesian decision-making framework, which are combined with incoming sensory information to form Bayes-optimal percepts of motion speed, orientation, convexity, and so on (W. J. Adams et al., 2004; Girshick et al., 2011; Peters et al., 2015; Stocker & Simoncelli, 2006). It has also been shown that internalization of natural scene statistics through experience can drive spontaneous neural activity in sensory areas, providing a neural basis for use of these statistics as a Bayesian prior (Berkes et al., 2011; Fiser et al., 2010).

We extend this idea to suppose that the system is sensitive to internal ‘natural scene statistics’ or empirical priors about *its own noise*, i.e. that the visual system has learned typical distributions of internal signal noisiness as a function of a stimulus’ location in the visual field. The system has likely learned that peripheral vision is noisier than central vision, forming the basis for an empirical prior over the reliability of internal representations of information sourced from these differing parts of the visual field. However, while such empirical priors are based in fact, they can sometimes lead to inaccurate perceptual estimates when the system is required to use them in Bayes-optimal computations. For example, empirical priors about man-made versus natural objects’ densities are used by the system in predicting visuo-haptic weight estimates (Peters et al., 2015), but in ways that lead to illusory percepts of heaviness due to simplification of complex, continuous priors over density into mixtures of Gaussians (Peters et al., 2016, 2018) -- perhaps due to biological restrictions on information coding (Heng et al., 2020). It has also been shown that highly asymmetric (skewed) priors over stimulus location which are initially learned correctly can be relied upon as if they were Gaussian when the observer is required to use them in a cue combination task (Acerbi et al., n.d.), and that incorrect priors in general can lead to illusions across many areas of visual perception in both healthy and atypical perception (Flanagan et al., 2008; Geisler & Kersten, 2002; Teufel et al., 2013; Valton et al., 2019; Weiss et al., 2002). Thus, incorrect priors can play a large role in ultimate percepts, even in first-order visual tasks.

We therefore reasoned that inflation of confidence in the visual periphery might likewise occur due to incorrect empirical priors about *noise* in the visual system as a function of visual field location, or at least incorrect usage of such priors. Specifically, we hypothesized that the true distribution of visual noise (variance or standard deviation of sensory signals) in the visual periphery is significantly positively skewed relative to central noise, due to mathematical properties of variance (i.e., it cannot be less than zero) and observation that peripheral vision is on average noisier (less reliable) than central vision. However, the metacognitive or even perceptual system may not be able to use this true prior to optimally estimate momentary noise levels, instead using a simplified, non-skewed prior for Bayesian noise estimation or even simply the mode of the prior, i.e. the most likely noise level to be encountered in the periphery. If the visual system uses incorrect priors, or otherwise incorrectly estimated the differences in reliability between central versus peripheral vision, such suboptimal introspection (Adler & Ma, 2018; Denison et al., 2018; Peters, Fesi, et al., 2017) could lead to variance misperception (Zylberberg et al., 2014) and thus over- as well as underconfidence.

In this project, we used five different Bayesian-inspired model observers to systematically explore the potential consequences of such a process. Our findings demonstrate how such an environmental statistic coupled with certain metacognitive errors in estimation or usage of this prior -- even under near-optimal Bayesian inference about noise -- could explain peripheral subjective inflation, especially under inattention.

## B. METHODS

### B.1. Conceptual description of models

We formalized the above intuition about the relative impact of different natural statistics of noise in central versus peripheral vision using signal detection & Bayesian decision theory (SDT/BDT) (Figure 1a).

**Figure 1.**
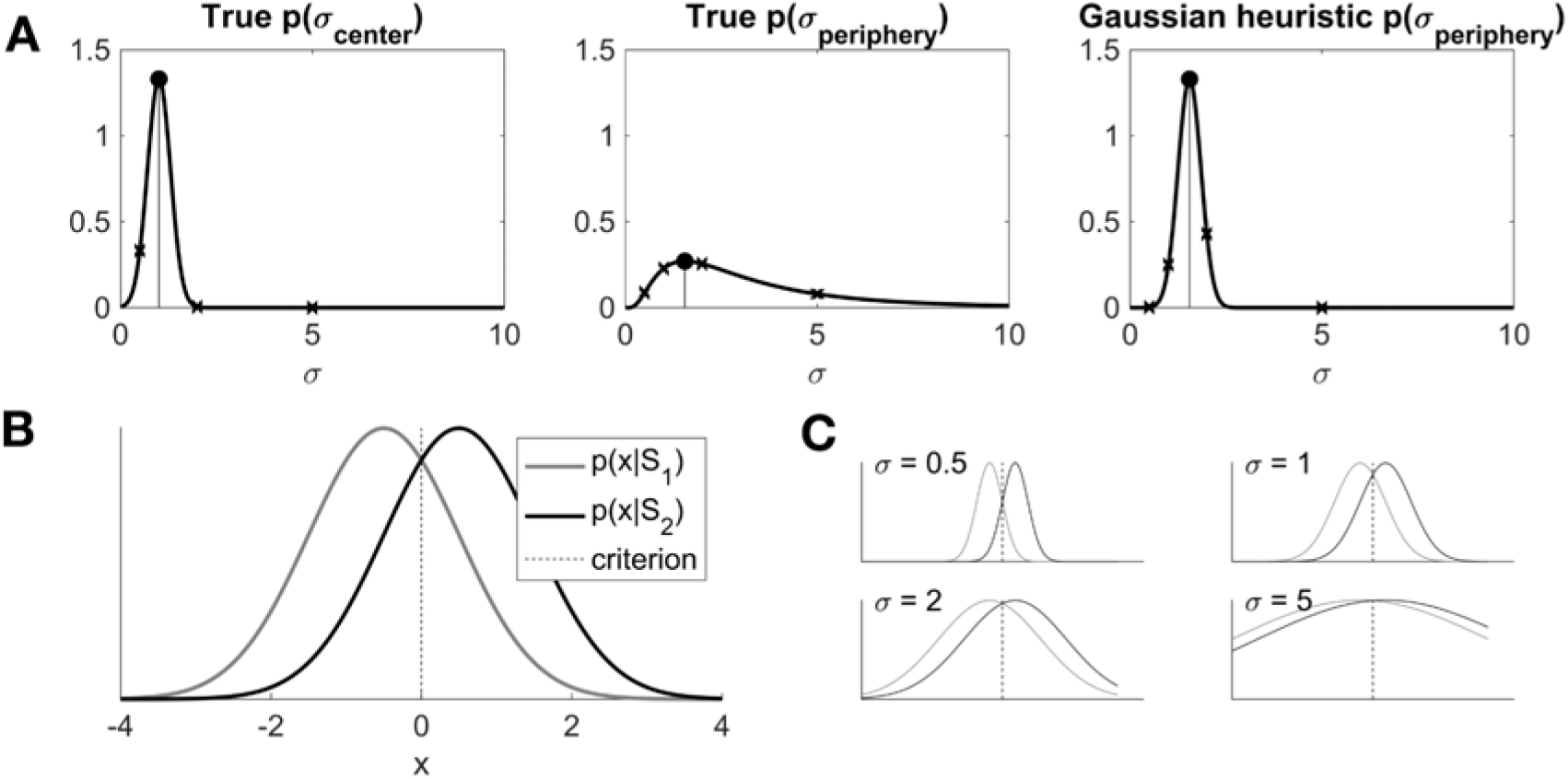
Hypothesized empirical prior distributions of noise in central and peripheral vision, and how they govern Type 1 decision-making in a Bayesian decision theory framework. (A) The empirical distribution of noise in central vision (left panel) is assumed to be Gaussian with relatively small mean and variance. The empirical distribution of noise in the visual periphery (middle panel) is assumed to be positively skewed with higher mean/median than in central vision. A heuristic metacognitive observer (e.g. model M5: hierarchical ‘Gaussian assumption’ heuristic observer; see Methods) may correctly represent the location and possibly scale of the peripheral noise distribution, but misrepresent its skewed shape as symmetric (right panel). Small x’s refer to possible noise levels sampled from these distributions, with consequences on the decision framework noted in (C).) (B) The noise in the central or peripheral visual field dictates the behavior of the Type 1 decision system, here represented in a signal detection or Bayesian decision theoretic framework. (C) As noise changes due to environmental conditions or location in the visual field (small x’s in the panels in (A)), the Type 1 decision system will become more or less precise, i.e. more or less sensitive to the signal in the environment.

#### B.1.i Type 1 decisions

We simulated an observer that engages in a simple discrimination task. On each trial, an observer samples the environment. The sample the observer receives may have come from a stimulus that has identity *S_1_*, or one with identity *S_2_*. Assuming Gaussian internal noise, these samples form Gaussian internal response distributions centered at the true mean for *S_1_* and *S_2_* with standard deviations governed by the combined internal noise of the observer and the external noise of the stimulus. For simplicity here we assume equal variances between the *S_1_* and *S_2_* distributions, following standard convention (Green & Swets, 1966; Macmillan & Creelman, 2005).

We modeled the distributions of these internal response signals as two Gaussian distributions representing cases where the signal came from *S_1_* or *S_2_* following standard SDT conventions (Green & Swets, 1966; Macmillan & Creelman, 2005) (Figure 1b). The discrimination criterion, which is hardcoded at the optimum in this simple system (halfway between the distributions to maximize performance, i.e. by setting the criterion where a sample is equally likely to have been generated by either stimulus class, which under equal priors corresponds to the location where the distributions intersect), is shown through a dashed vertical line. When the internal response variable falls above this threshold, the observer answers “*S_2_*” to a discrimination question, and otherwise answers “*S_1_*”. The internal response criterion in the models below is specified in posterior probability ratio space, making this observer a Bayesian observer, to facilitate the confidence readout as the posterior probability of the choice that the observer made. See Section B.2 Formal Computational Models for specific details.

To this simple Type 1 system we add an additional hierarchical or metacognitive (Type 2) inference layer to explore several hypotheses about how the noise in the system (both internal noise and external noise (Lu & Dosher, 2008)) governs the Type 1 decision space. Drawing inspiration from (1) Bayesian ideal observer analysis demonstrating that natural scene statistics govern Type 1 perception (W. J. Adams et al., 2004; Girshick et al., 2011; Peters et al., 2015; Stocker & Simoncelli, 2006), and (2) hierarchical models in which a Bayesian ideal observer’s inference about latent variables also governs the ultimate percept (Knill & Richards, 1996; Knill & Saunders, 2003; Körding et al., 2007; Körding & Tenenbaum, 2007a; Odegaard et al., 2015; Peters et al., 2016, 2018; Samad et al., 2015; Yuille & Bülthoff, 1996), we developed and compared a series of ‘flat’ and ‘hierarchical’ inference models with varying ‘knowledge’ or reliance of natural scene statistics of noise in central versus peripheral vision to evaluate their predictions for peripheral inflation.

#### B.1.ii Environmental statistics of noise in central versus peripheral vision

A critical factor in an observer’s ability to make accurate Type 1 decisions is the noisiness (variability) of the system itself: noise can cause the observer to respond “*S_1_*” when a signal actually came from the *S_2_* distribution, or vice versa, with varying levels of precision or error (Figure 1d). The noise in the system depends on the location in the visual field, central or peripheral, with central vision being more reliable (less noisy) than peripheral vision (Provis et al., 2013); (Gloriani & Schütz, 2019). It is therefore reasonable to assume that central and peripheral vision have different hypothesized underlying (parent) distributions governing the distributions (Lu & Dosher, 1998) of noise typically experienced in these two regions of the visual field (Figure 1b & c). Specifically, the variance (noise) corresponding to peripheral vision is assumed to be higher than for central vision.

A Bayesian observer that is sensitive to environmental statistics about a given variable will represent expectations about such variables as prior distributions -- for example, about light source location, motion speed, or contour orientation (W. J. Adams et al., 2004; Girshick et al., 2011; Stocker & Simoncelli, 2006), as mentioned above. Here, we hypothesized that central and peripheral environmental distributions of *noise* experienced by the visual system also lead the visual system to form prior expectations about variability as a latent variable, following previous convention in hierarchical Bayesian inference in vision and multisensory integration (Beierholm et al., 2009; Knill & Richards, 1996; Knill & Saunders, 2003; Körding et al., 2007; Körding & Tenenbaum, 2007a; Landy et al., 2011; Odegaard et al., 2015; Peters et al., 2015, 2016, 2018; Samad et al., 2015; Shams et al., 2000; Wozny et al., 2008; Yuille & Bülthoff, 1996). That is, the visual system learns to expect that central vision typically involves less noisy signals, while peripheral vision typically involves noisier signals.

Our hypothesis is further extended by the observation that variance is a property that cannot be mathematically negative, such that any distribution of variance must by definition exist in the domain *σ* > 0. When the distribution of variance is narrow and symmetric, a truncated Gaussian distribution with small mean (and with domain of *σ* > 0) might be modestly appropriate to represent the empirical distribution for central noise (Figure 1a); that is, we hypothesized that the empirical distribution of noise is appropriately represented by a *symmetric* distribution around a typical (mode/most common) level of noise, which for central vision is rather small. However, in the periphery it is possible that a broader distribution of noise is experienced, especially under varied lighting conditions, and that this distribution ought also to be across higher noise values in general. Although it would be possible for such a distribution to be symmetric as in central vision, here we hypothesized a positively skewed empirical distribution for noise in the periphery (Figure 1b); that is, we hypothesized that when the variance does fluctuate it does so *asymmetrically* around a typical (mode/most common) level of (typically higher than central) noise. We note that although such a distribution is mathematically justified by properties of noise (variance), it has yet to be empirically demonstrated (see Discussion). Thus, the purpose of this project was to serve as a precursor to measuring such statistics, and their potential use by observers, by evaluating the impact of such natural statistics on decisions and confidence judgments -- including how an observer’s differential ‘knowledge’ of this true empirical distribution might explain puzzling behaviors such as peripheral subjective inflation.

#### B.1.iii Intuitive introduction to models

Typically, the visual system is assumed to be optimal (Bejjanki et al., 2016; Landy et al., 2007) in that it uses an accurate estimate of its own noise because it has ‘knowledge’ of its own internal statistics (King & Dehaene, 2014; Lau, 2008); this optimality is assumed to propagate to the visual metacognitive system (Drugowitsch, 2016; Drugowitsch et al., 2019; Fleming & Daw, 2017), perhaps corrupted by some additional metacognitive noise (Maniscalco & Lau, 2016). However, a number of heuristic models have previously been shown to capture metacognitive behavior better than optimal models, including variance misperception (Peters, Fesi, et al., 2017; Zylberberg et al., 2014) (this is also seen in cognitive decision-making; (Herce Castañón et al., 2019)), fixed Type 2 (metacognitive) criterion setting under varying noise (Li et al., 2018; Rahnev et al., 2011; Rahnev, Maniscalco, et al., 2012; Solovey et al., 2014), a bias indicating overweighting evidence favoring a decision (Maniscalco et al., 2016; Peters, Thesen, et al., 2017; Zylberberg et al., 2012), or suboptimal Type 2 criterion setting (Adler & Ma, 2018), among others. In particular, overconfidence relative to true Type 1 sensitivity (subjective inflation; also related to detection biases) is more often observed in the visual periphery than the center (Li et al., 2018; Odegaard et al., 2018; Rahnev et al., 2011; Solovey et al., 2014)(Li et al., 2018; Rahnev et al., 2011; Solovey et al., 2014).

Combining these observations, we hypothesized that the visual system in both peripheral and central vision may not be able to accurately perceive the noisiness of the stimulus. To explore how various heuristic strategies could lead to overconfidence specific to the periphery, we defined and compared five candidate computational models, which differ according to whether and how they estimate or know about noise differently across central versus peripheral vision. Simple descriptions of each model are provided in Table 1; we provide full details of these models in Methods.

**Table 1.**
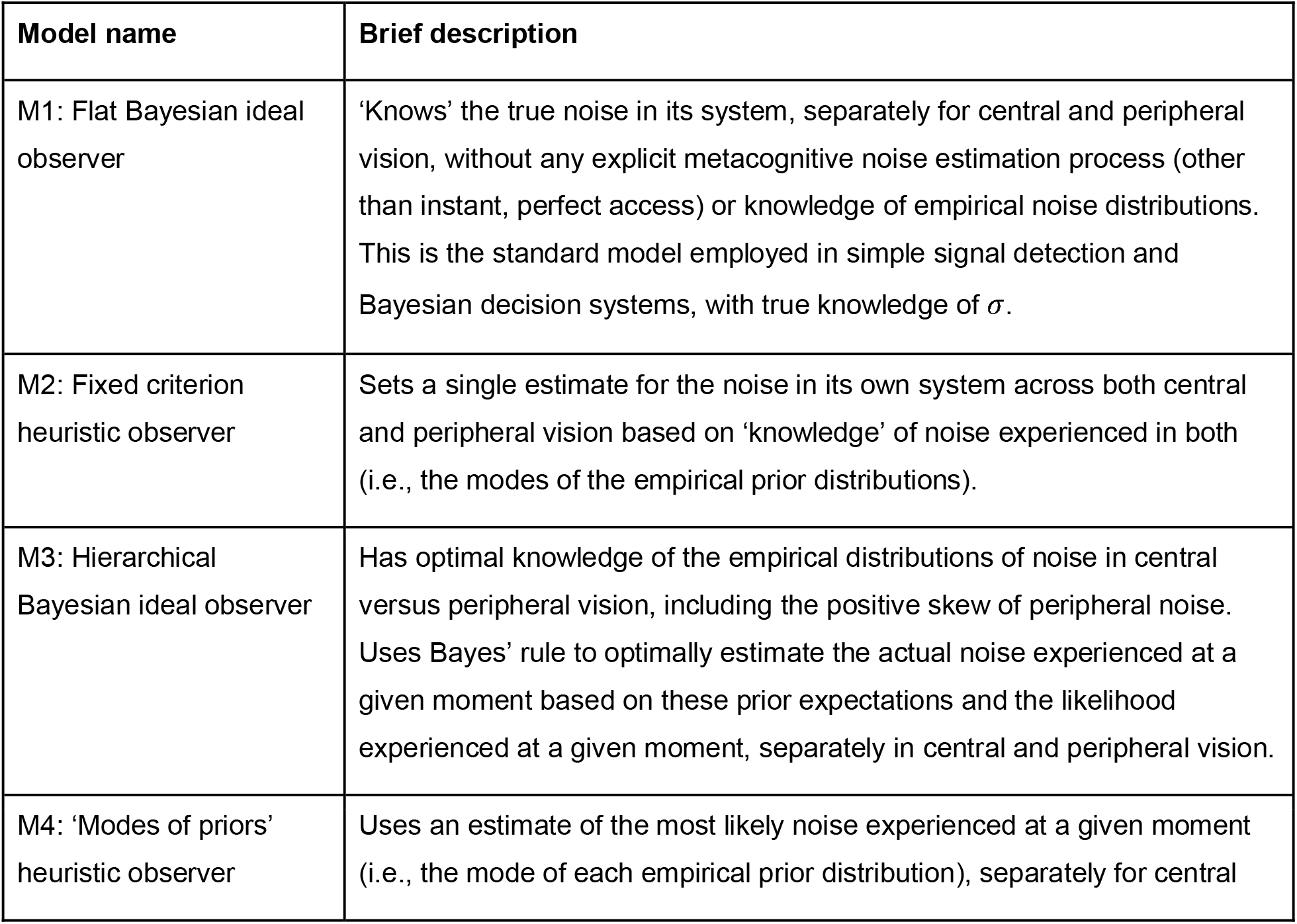
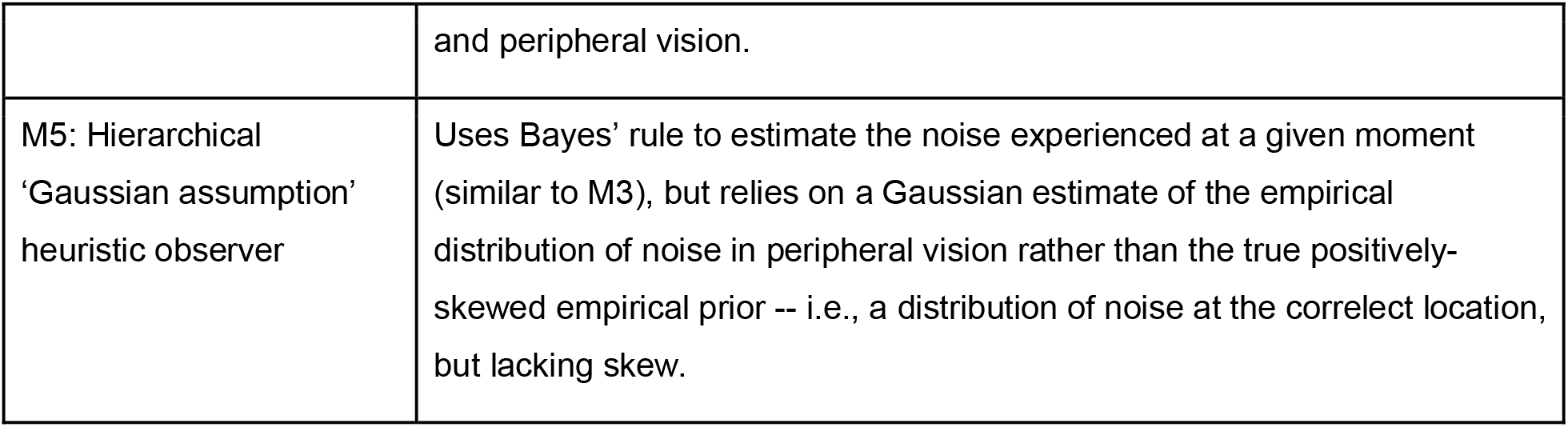
Introduction to the five Bayesian ideal and heuristic models tested.

| Model name | Brief description |
| --- | --- |
| M1: Flat Bayesian ideal observer | ‘Knows’ the true noise in its system, separately for central and peripheral vision, without any explicit metacognitive noise estimation process (other than instant, perfect access) or knowledge of empirical noise distributions. This is the standard model employed in simple signal detection and Bayesian decision systems, with true knowledge of $\sigma$. |
| M2: Fixed criterion heuristic observer | Sets a single estimate for the noise in its own system across both central and peripheral vision based on ‘knowledge’ of noise experienced in both (i.e., the modes of the empirical prior distributions). |
| M3: Hierarchical Bayesian ideal observer | Has optimal knowledge of the empirical distributions of noise in central versus peripheral vision, including the positive skew of peripheral noise. Uses Bayes’ rule to optimally estimate the actual noise experienced at a given moment based on these prior expectations and the likelihood experienced at a given moment, separately in central and peripheral vision. |
| M4: ‘Modes of priors’ heuristic observer | Uses an estimate of the most likely noise experienced at a given moment (i.e., the mode of each empirical prior distribution), separately for central |
|  | and peripheral vision. |
| M5: Hierarchical<br>'Gaussian assumption'<br>heuristic observer | Uses Bayes' rule to estimate the noise experienced at a given moment (similar to M3), but relies on a Gaussian estimate of the empirical distribution of noise in peripheral vision rather than the true positively-skewed empirical prior -- i.e., a distribution of noise at the correct location, but lacking skew. |

For each of these models, we estimated the behavior of the Type 1 and Type 2 decision-making system under different variances, i.e. different levels of noise in the signal, under central versus peripheral vision conditions. The critical factor to evaluate the models’ performance is how each model makes metacognitive decisions relative to Type 1 performance capacity (which is dictated by the true noise regardless of the metacognitive noise estimation process). Model details are presented in the next section, Section B.2 Formal Computational Models.

### B.2 Formal computational models

#### B.2.i Type 1 decisions

Two Gaussian distributions represent the internal response distributions for two signal sources, *S_1_* and *S_2_* (Figure 1b). To make a decision, the conditional probability of the source being present given an internal response variable on that trial is calculated through Bayes rule:

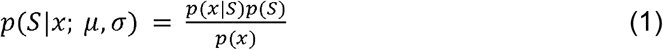

where *S* is source (*S_1_* or *S_2_*), and *x* is the internal response, with

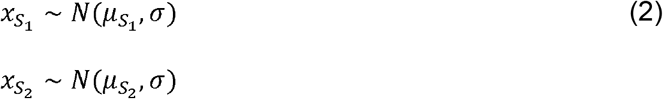

where 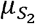 is the mean of the internal response to a signal from distribution S2 with a given strength, 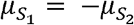 following previous convention, and *σ* represents their (assumed to be equal) standard deviations under the simplest implementation. Prior probabilities for sources *S_1_* and *S_2_* are defined by

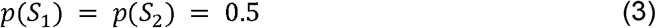

The decision made (D) is determined by the following

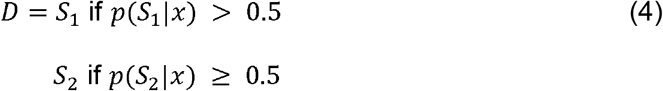

where 0.5 serves as the optimal criterion in an SDT framework (set to maximize percent correct choices). That is, the criterion is set so the observer selects *S_i_* according to which is most likely to be correct.

The standard convention is that confidence is computed via

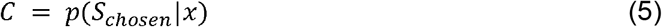

The models below selectively alter elements of this standard process, with focus on Equations 1 and 2.

#### B.2.i Empirical prior distributions governing noise in central versus peripheral vision

The standard deviations of both the 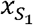 and 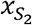 internal response distributions are set to the same *σ* (Equation 2) within a given visual field location, with *σ* governed by the noise distributions at each location: central vision’s empirical distribution of standard deviations (i.e., the natural statistics of noise in central vision) is symmetric (Gaussian), while peripheral vision’s is positively skewed:

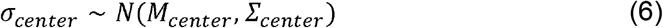

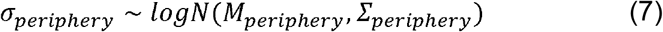

where *M_center_* and *M_periphery_* represent the mean (or log mean) of true standard deviations experienced by central and peripheral visual fields across a wide range of situations, respectively, *Σ_center_* and *Σ_periphery_* represent the respective variabilities in these empirical distributions, and *σ* is always constrained to be in domain *σ* > 0 (i.e., Equation 6 is a truncated normal distribution). Thus, across all simulated conditions, central vision experiences a symmetric distribution of noise around a central mean (Figure 1a), while peripheral vision experiences on average higher noise and with positive skew (Figure 1b).

#### B.2.ii Definitions of Bayesian ideal and heuristic models of metacognition

##### M1: Flat Bayesian ideal observer

This observer ‘knows’ the true noise in its system, separately for central and peripheral vision, without any explicit metacognitive noise estimation process or knowledge of empirical noise distributions. This is the standard model employed in simple signal detection and Bayesian decision systems, with true knowledge of *σ*. This model calculates the Type 1 decision via Equations 1, 2 and 4, and confidence via Equation 5 as written above.

##### M2: Fixed criterion heuristic observer

This observer sets a single estimate for the noise in the environment across both central and peripheral vision based on optimal ‘knowledge’ of both empirical noise distributions. Thus, it is conceptually akin to the ‘fixed Type 2 criterion’ model described in previous literature (Li et al., 2018; Rahnev et al., 2011; Solovey et al., 2014). This model alters Equations 1 & 2 such that all decisions and confidence judgments for both central and peripheral vision are based on a single heuristic estimate of *σ*, 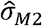, which can be thought of as halfway between the peaks of the two noise prior distributions for center and periphery, i.e. an “average amount of noise”:

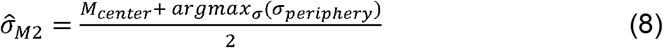

Thus, Equation 1 is rewritten as:

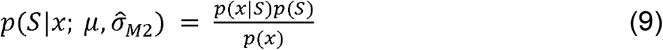

with Equation 2 rewritten as the observer *assuming*

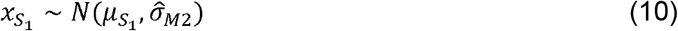

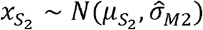

Decisions and confidence judgments are made according to Equations 4 and 5 as above.

##### M3: Hierarchical Bayesian ideal observer

This observer has optimal knowledge of the empirical distributions of noise in central versus peripheral vision, including the positive skew of peripheral noise. It uses Bayes’ rule to optimally estimate the actual noise experienced at a given moment based on these prior expectations and the likelihood experienced at a given moment, separately in central and peripheral vision. That is, this model alters Equations 1 & 2 such that all decisions and confidence judgments for both central and peripheral vision are based on a optimal estimates of *σ*, 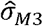, with one estimate for each of center and periphery:

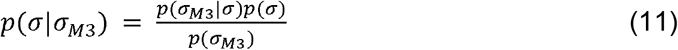

with assumed Gaussian noise in estimating self-noise, i.e. *p*(*σ*_*M*3_|*σ*) follows a normal distribution with mean *σ* (the noise present in the stimulus and system in this particular condition or trial) and standard deviation of this likelihood distribution defined as *ς* -- again with separate definitions *ς* for each of the (separable) center and periphery. *p*(*σ*)refers to the empirical prior distribution of noise for a given visual field location (center or periphery). That is, there is an amount of noise that is currently happening, but the observer does not have ‘perfect’ access to this noise; instead, it has a noisy representation of this noise (akin to how one would have a noisy representation of some other aspect of a physical stimulus [length, location, size, speed, etc.] which is combined with prior expectations for that noise (Alais & Burr, 2004; Beierholm et al., 2009; Burge & Girshick, 2010; Girshick et al., 2011; Knill, 2003, 2007; Knill & Richards, 1996; Knill & Saunders, 2003; Körding et al., 2007; Landy et al., 2011; Odegaard et al., 2015; Peters et al., 2018; Weiss et al., 2002; Wozny et al., 2008, 2010; Yuille & Bülthoff, 1996). Note that this suggests that within a given trial, the metacognitive system may have access to more than one sample of noise, which are normally distributed around the true noise being experienced according to *p*(*σ*_*M*3_|*σ*) ~ *N*(*σ*, *ς*) (although we do not explicitly model this metacognitive sampling process; see below for simulation details). The estimate for 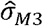 is chosen as the maximum a posteriori estimate, i.e.

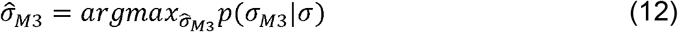

Subsequently, decisions and confidence are made as above by altering Equations 1 & 2 to read for each of the central and peripheral judgments:

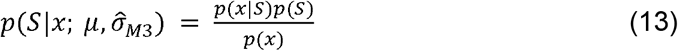

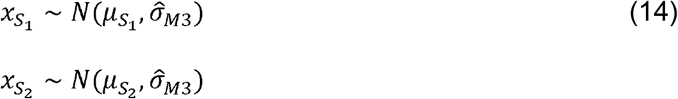

and then following Equations 4 and 5 as above to complete the readout.

##### M4: ‘Modes of priors’ heuristic observer

This observer uses an estimate of the most likely noise experienced at a given moment, but unlike M2 does this separately for central and peripheral vision. Rather than optimally estimating (or just knowing) the noise as the M3 or M1 observers do, it instead uses the mode of each empirical prior distribution as a heuristic estimate -- it picks the most likely amount of noise experienced at this location in the visual field. That is, rather than using *σ*, this observer sets 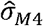 via

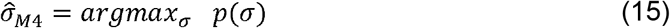

again, separately for center and periphery as before. Then, just as M3, we redefine decisions and confidence by rewriting Equations 1 & 2 to reflect these assumptions for each of the central and peripheral judgments:

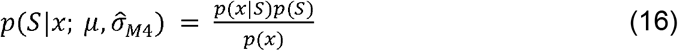

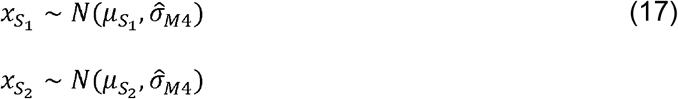

and then following Equations 4 and 5 as above to complete the readout.

##### M5: Hierarchical ‘Gaussian assumption’ heuristic observer

This observer uses Bayes’ rule to estimate the noise experienced at a given moment (similar to M3), but assumes that the empirical distribution of noise in peripheral vision is Gaussian rather than using the true positively-skewed empirical prior. (It also uses a Gaussian assumption for central vision, but since this is accurate, the central vision M5 model is identical to the central vision M3 model.) We assume that the observer understands the general location (magnitude) of this peripheral noise distribution, but possesses poorer knowledge of its variability or level of skewness.

Thus, this observer first estimates the location (mode) of the empirical prior governing peripheral noise as:

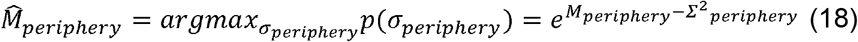

which follows from the fact that the peripheral noise distribution is lognormal (Equation 7). That is, the location (mean and mode) of the “estimated”, symmetrical prior for *σ* in the visual periphery is computed so as to match the most probable noise level (mode) of the “true,” asymmetrical prior.

Variability in the estimated peripheral noise empirical prior distribution is then set to some multiple of the variability in the central noise prior:

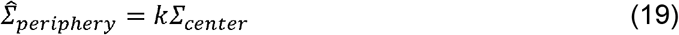

The strength of this assumption (i.e., the magnitude of *k*, the variability assumed in the peripheral noise priori) controls the strength of peripheral inflation but does not qualitatively affect results. Because the purpose of the present project is to provide a proof of concept, we do not fit *k* here, and set *k* = 1 for simplicity. Future work ought to fit *k* to empirical data.

Then, we redefine the *estimated* empirical prior for noise in the periphery (Equation 7) for use by this observer as:

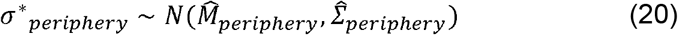

(The corresponding distribution for *σ*\*_*center*_ is equivalent to *σ*_*center*_, since 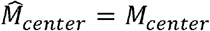 and 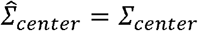 due to the correct expectation Gaussian noise distributions in the visual center.) Then, just as with M3, all decisions and confidence judgments for both central and peripheral vision are based on a optimal estimates of *σ* (now based on the incorrect prior expectations for the distribution of *σ*\*), 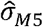, again for each of center and periphery:

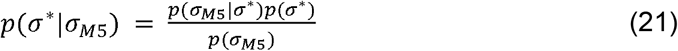

with assumed Gaussian noise in estimating self-noise, i.e. *p*(*σ*_*M*5_|*σ*\*) follows a normal distribution with mean *σ*\* and standard deviation *ς*, again for each of the center and periphery. That is, *p*(*σ*\*)now refers to the *incorrect* expected prior distribution of noise for a given visual field location (center or periphery). As with M3, the estimate for 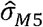 is chosen as the maximum a posteriori estimate, i.e.

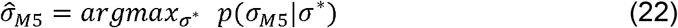

Subsequently, as with M3, decisions and confidence are made by altering Equations 1 & 2 to reflect these assumptions, reading for each of the central and peripheral judgments:

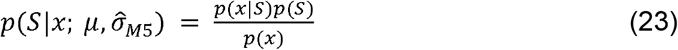

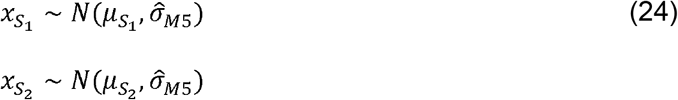

and then following Equations 4 and 5 as above to complete the readout.

### B.3 Attentional manipulations

Subjective inflation of confidence relative to performance has been shown previously to be stronger under cases of inattention (Li et al., 2018; Rahnev et al., 2011), such that -- somewhat counterintuitively -- increased sensory precision due to attention leads to a reduction in peripheral subjective inflation in particular. We next investigated the behavior of all five model observers under simulated conditions of increasing attention by assuming that attention may modify two factors: (1) the precision in the sensory estimate (i.e., the means of the empirical distributions of noise, *M*), and (2) the precision in the metacognitive estimate of noise (i.e., the standard deviations of the likelihood distributions of noise, *ς*). For this simple proof of concept, we introduced two additional scalar multiplicative factors that could modify these two parameters for both Type 1 and Type 2 (metacognitive) judgments: *a*_1_ and *a*_2_ for each noise factor, respectively. That is, *M* is actually defined as *a*_1_*M* and *ς* as *a*_2_*ς*, with *a*_1_ = *a*_2_ =1 under default (‘neutral’) conditions. Thus, under increasing attention, *a*_1_ and *a*_2_ are both assumed to decrease, increasing the precision of the Type 1 samples themselves as well as the Type 2 (metacognitive) estimates about noise. Increasing attention is referred to as ‘attention+’ (*a*_1_ = 0.9, *a*_2_ = 0.8) and ‘attention++’ (*a*_1_ = 0.8, *a*_2_ = 0.6) with reference to ‘neutral’ (no attentional manipulation) in Results.

Finally, to provide a more quantitative assessment of how models behave under the specific performance (*d’*) and attentional manipulation magnitudes reported in the empirical literature, we conducted qualitative fitting of Odegaard et al. (2018)’s Experiment 1. This study used crowding as an attentional manipulation to manipulate performance and confidence in the visual periphery. It is notable that the majority of studies in this area typically ask about visibility or detection, which -- while conceptually similar -- is not the same thing (Sandberg et al., 2010; Wierzchoń et al., 2014)he target of our models. Thus, we opted to conduct qualitative fitting to the summary statistics reported by Odegaard and colleagues. They found that in the unattended (crowded) condition, Type 1 performance (*d’*) was reduced from about 1.5 to 1.1 (their Figure 2); however, while confidence for correct trials did not change between attended and unattended conditions, confidence for incorrect trials *increased* in unattended trials (their Figure 3). To capture this result, we first expanded our simulation to include a larger range of “signal strength” levels to vary between 1 ≤ *μ*_*S*2_ ≤ 10 (in the main simulations, it is set simply to μ_S2_ = 0 and μ_S2_ = 1; Table 1). Next, we selected combinations of signal strength and noise level from this expanded simulation that produced d’ values of approximately 1.5 (between 1.4-1.6) in our “attention++” condition, and approximately 1.1 (between 1.0 and 1.2) in our “neutral” condition. All other parameters were held constant from the main simulations.

**Figure 2.**
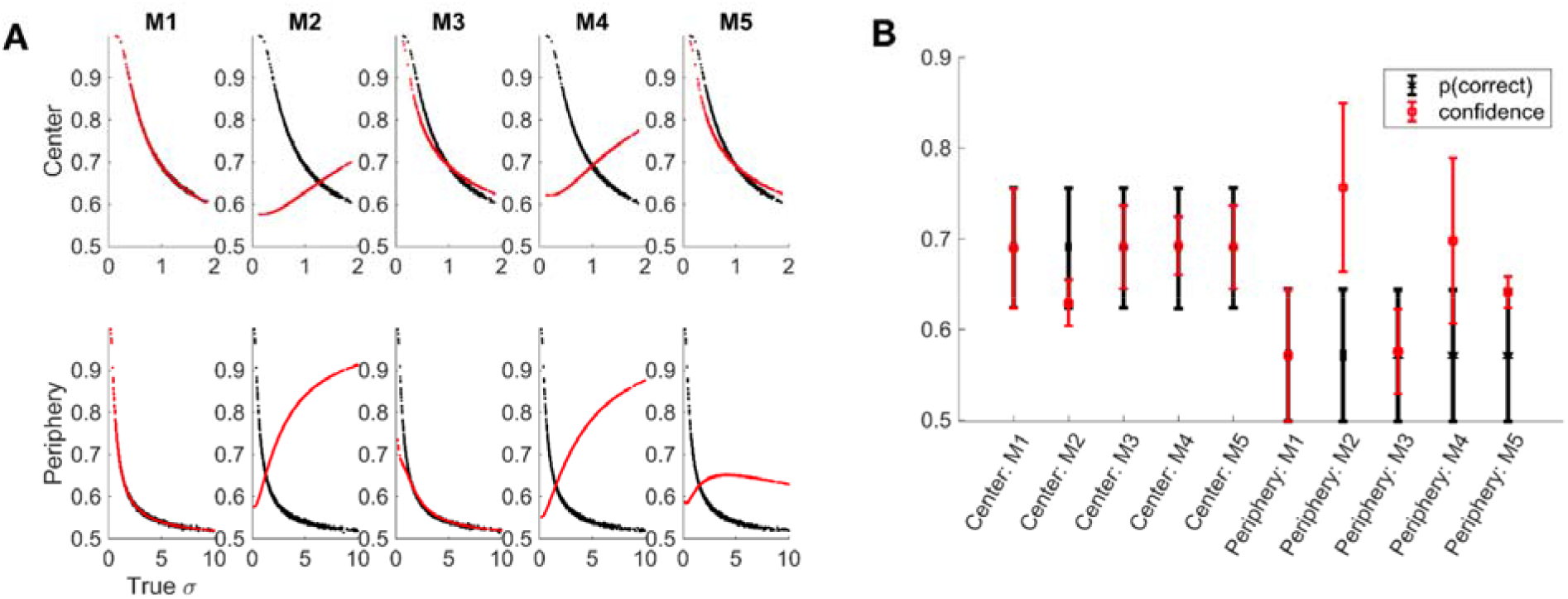
Predicted performance and confidence behavior from simulations of all five model observers. (A) Predicted performance (p(correct)) and confidence in central and peripheral vision for each model across a range of noise levels. (B) Median performance and confidence for each model observer in central and peripheral vision. Models M1 (flat Bayesian ideal observer) and M3 (hierarchical Bayesian ideal observer) display close match between performance (p(correct)) and confidence in both central and peripheral vision, leading to neither under- nor overconfidence on average. In contrast, models M2 (fixed criterion heuristic observer) and M4 (‘modes of priors’ heuristic observer) display patterns of both under- and overconfidence in central and peripheral vision. Finally, while M5 (hierarchical ‘Gaussian assumption’ heuristic observer) correctly estimates confidence in central vision due to knowledge of the shape of the empirical prior distribution of noise, due to its assumption of symmetry in the visual periphery noise distribution, it systematically under-estimates noise leading to average over-confidence for the visual periphery only. See Table 1 and Methods for details of models.

**Figure 3.**
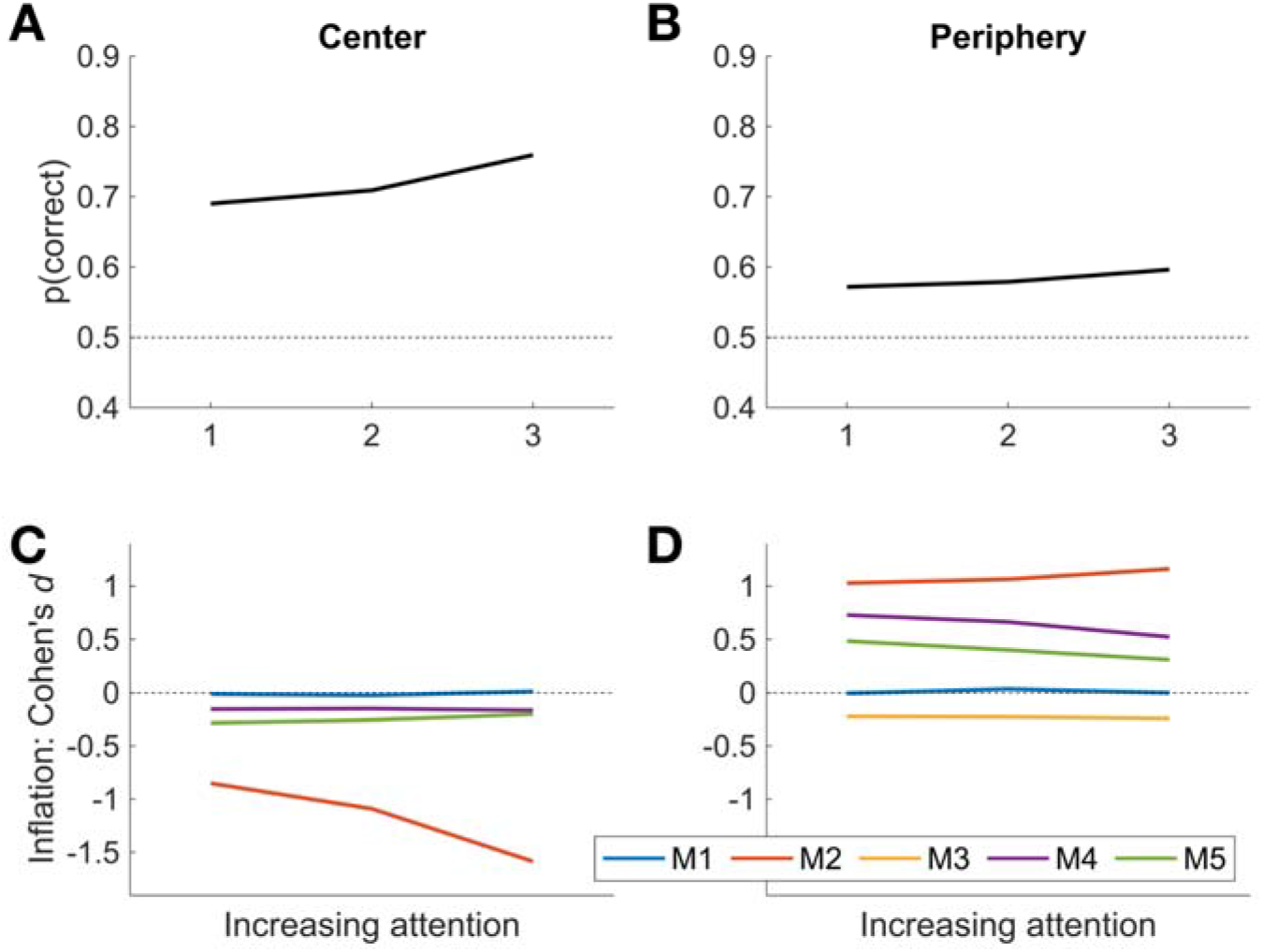
Results of simulated attentional manipulations on the effect size (Cohen’s *d*) of subjective inflation (overconfidence). Increasing attention is modeled as increasing precision in sensory estimates and increasing metacognitive noise estimation precision (see Methods). In central vision, increasing attentional allocation leads as expected to improved Type 1 performance (A), but generally flat or slightly increasing effect size for inflation (C) with the exception of model M2, which shows stronger and stronger *under*confidence as attention increases. In contrast, in the periphery, despite attentional allocation increasing performance as expected (B), most models display flat or slightly increasing effect size for peripheral inflation (D) with the exception of models M4 and M5, the two heuristic hierarchical models.

### B.4 Simulation details

We used Monte Carlo simulations to evaluate the performance and confidence behavior of all five models described above, assuming the same signal strength in all scenarios for the main simulations. Parameters for all simulations are presented in Table 2; although these parameter values are arbitrary for the purposes of demonstration, the models’ qualitative behavior is similar across a range of parameter values (data not shown).

**Table 2.** Parameters used in the main simulations. *Note that for the simulations to fit data reported by Odegaard and colleagues (2018), we expanded the range for the mean of *S_2_* to be 1 ≤ *μ*_*S*2_ ≤ 10; all other parameters for this additional fit remained the same as for the main simulations.

| Parameter | Description | Used by | Value(s) |
| --- | --- | --- | --- |
| $\mu_{S_1}$ | mean of $S_1$ distribution | all models | 0 |
| $\mu_{S_2}$ | mean of $S_2$ distribution | all models | 1* |
| $M_{center}$ | mean of central vision noise prior distribution | M3, M4, M5 | 1 |
| $M_{periphery}$ | mean of peripheral vision noise prior distribution | M3, M4, M5 | 1 |
| $\Sigma_{center}$ | standard deviation of central vision noise prior distribution | M3, M4, M5 | 0.3 |
| $\Sigma_{periphery}$ | standard deviation of peripheral noise prior distribution | M3, M4, M5 | 0.75 |
| $\varsigma_{center}$ | standard deviation of central noise likelihood distribution | M3, M5 | 0.15 |
| $\varsigma_{periphery}$ | standard deviation of peripheral noise likelihood distribution | M3, M5 | 0.5 |
| $a_1$ | scaling factor for the mean of the noise prior distributions | all models | [1,0.9,0.8] |
| $a_2$ | scaling factor for the standard deviation of the noise likelihood distributions | M3, M5 | [1,0.8,0.6] |

For each of the above-described model observers (M1 - M5), we sampled 1000 standard deviations *σ* from each of the central and peripheral distributions (samples with *σ* < 0 were set to *σ* = *eps* =2.2204e-16); at each sampled standard deviation we calculated the performance (% correct) and average confidence (mean *C*) for each model across 50,000 simulated trials (25,000 *S_1_* and 25,000 *S_2_*). Note that this means that while noise varies across conditions, it is not assumed to vary across trials within a condition. We quantified over- and underconfidence using Cohen’s *d* as a measure of effect size, with positive values indicating confidence > performance and negative values indicating confidence < performance:

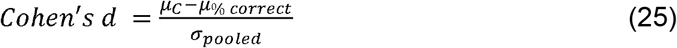

with *μ_C_* the mean of confidence judgments (Equation 5), *μ*_% *correct*_ the proportion of correct responses, and *σ*_*pooled*_ their pooled standard deviation (for paired samples) according to standard definitions. All simulations were carried out through custom scripts written in Matlab R2019b.

## C. RESULTS

### C.1 Central versus peripheral confidence relative to performance (no attentional manipulation)

We first examined the behavior of the five simulated observers under ‘default’ conditions, i.e. using the parameters presented in Table 1 without any attentional manipulations (‘neutral’, *a*_1_ = *a*_2_ = 1). The key measure is that of confidence inflation, measured here using the effect size measure of Cohen’s *d*: mean difference between confidence (Equation 5) and the percent correct answers (performance governed by Equation 4), scaled by their pooled standard deviation, such that positive values of *d* indicate confidence > performance, while and negative values indicate confidence < performance. See Methods for details.

#### C.1.i Model M1: Flat Bayesian ideal observer

The flat Bayesian ideal observer (M1) followed optimal expected behavior in both central and peripheral vision (Figure 2a, first column). Under higher noise conditions performance (p(correct)) falls and this drop is mirrored in confidence estimates for both visual field locations. Thus, this observer displayed neither under- nor overconfidence on average (Figure 2b). The average effect size for M1 in the center was Cohen’s *d* = −0.0104, and in the periphery was Cohen’s *d* = −0.0054 -- that is, there is almost no overconfidence on average for either visual field.

#### C.1.ii Model M2: Fixed criterion heuristic observer

The fixed criterion heuristic observer (M2) also behaved as reported in previous literature (Li et al., 2018; Rahnev et al., 2011; Solovey et al., 2014) (Figure 2a, second column). Performance dropped as expected for increasingly noisy conditions in both central and peripheral vision, but due to the fixed Type 2 criterion (set here according to an average of the noise in central and peripheral visual fields) M2 displayed a general underconfidence bias under less noisy conditions and a general overconfidence bias under noisier conditions. Importantly, because peripheral vision is on average much noisier than central vision, the fixed confidence criterion led to biases towards underconfidence relative to performance in central vision, accompanied by significant peripheral inflation (overconfidence relative to performance) (Figure 2b). The average effect size for M2 in the center was Cohen’s *d* = −0.8510 (i.e., strong underconfidence), and in the periphery was Cohen’s *d* = 1.0273 (i.e., strong overconfidence). If the fixed criterion were instead set only according to central vision noise, this would translate to optimal confidence in central vision and extreme overconfidence in the periphery; however, this is not the focus of the present project, so we leave such explorations to future studies.

#### C.1.iii Model M3: Hierarchical Bayesian ideal observer

The hierarchical Bayesian ideal observer (M3) estimates the noise in its system optimally using both prior experience (optimal knowledge of the empirical prior governing noise in central versus peripheral vision) and an estimate of noise at the given moment. Thus, it also displayed similar behavior to model M1, albeit with slight bias towards underconfidence under extremely not-noisy conditions and towards overconfidence under extremely noisy conditions due to reliance on knowledge of the empirical distributions of noise (i.e., use of the prior distributions of noise) (Figure 2a, third column). Despite these slight biases, M3 displayed neither under- nor overconfidence on average in either central or peripheral vision (Figure 2b). The average effect size for M3 in the center was Cohen’s *d* = −0.2832, and in the periphery was Cohen’s *d* = - 0.2212 -- that is, both central and peripheral vision displayed modest underconfidence, but in similar magnitude.

#### C.1.iv Model M4: ‘Modes of priors’ heuristic observer

The ‘modes of priors’ heuristic observer (M4) uses the most probable level of noise in each of the central and peripheral visual fields, based on its previous experience, rather than represent and use the entire empirical prior (as in M3) or use a single estimate across both center and periphery (as in M2). Due to the fact that this empirical distribution is (here hypothesized to be) symmetrical around a typical noise level in central vision, this strategy leads M4 to exhibit on average no over- or underconfidence in the center of the visual field (Figure 2a, fourth column, top row; Figure 2b). In contrast, like M1 the M4 observer displayed overconfidence in the visual periphery -- but this time, it is due to the fact that the peripheral noise distribution is (here hypothesized to be) positively skewed. Thus, reliance on the mode of the empirical prior distribution led to overconfidence more often than underconfidence, and overconfidence on average in the visual periphery (Figure 2a, fourth column, bottom row; Figure 2b). The average effect size for M4 in the center was Cohen’s *d* = −0.1522 (i.e., modest underconfidence) and in the periphery was Cohen’s *d* = 0.7287 (i.e., strong overconfidence).

#### C.1.v Model M5: Hierarchical ‘Gaussian assumption’ heuristic observer

Finally, we examined the predicted behavior of the hierarchical ‘Gaussian assumption’ heuristic observer (M5). M5 is nearly identical to M3 (the hierarchical Bayesian ideal observer) with the important difference that M5 uses a Gaussian estimate for the empirical distribution of noise in the visual periphery. Thus, while M5 displayed identical behavior to M3 in the center (Figure 2a, fifth column, top row; Figure 2b), this Gaussian assumption for peripheral noise led M5 to more often under- than over-estimate noise in the visual periphery, leading once again to peripheral overconfidence on average (Figure 2a, fifth column, bottom row; Figure 2b). Thus, even though M5 possesses knowledge that the empirical distribution of noise in the visual periphery is generally much noisier than in the center, its assumption that this distribution is symmetrical led to an average under-estimation of noise and therefore overconfidence relative to performance. The average effect size for M5 in the center was Cohen’s *d* = −0.2857 (i.e., modest underconfidence because it is identical to M3), but in the periphery was Cohen’s *d* = 0.4846 (i.e., medium-strong overconfidence).

### C.2 Attentional manipulations

Given that several of the model observers we tested displayed overconfidence in the visual periphery, we next turned to attentional manipulations as a possible avenue for arbitrating models’ fit to previously-reported effects. In particular, it has previously been reported that attentional manipulations may affect peripheral inflation in a somewhat counterintuitive manner, with high-attention conditions leading to less strong peripheral overconfidence and inattention causing increases in subjective inflation in the periphery (Li et al., 2018; Rahnev et al., 2011; Solovey et al., 2014). Thus, we simulated anticipated effects of increasing attention as decreases in sensory noise and increases in metacognitive precision (see Methods). We quantified the effect size of inflation, i.e. the difference between performance and confidence, as Cohen’s *d*, with positive values indicating confidence > performance and negative values indicating confidence < performance.

As expected, increasing attentional allocation increased sensory precision and thus increased performance in both central and peripheral vision (Figure 3a,b; Table 3). However, this was generally accompanied by a relatively flat or if anything slightly increasing effect size (Cohen’s *d*) for subjective inflation (confidence overestimating performance) for both central and peripheral vision (Figure 3c,d; Table 3), with the exceptions of model M2 in central vision (Figure 3c; Table 3) and models M4 and M5 in peripheral vision (Figure 3d; Table 3).

**Table 3.** Cohen’s *d* measures of effect size for inflation, i.e. the difference between confidence (*p*(*choice*|*x*)) and performance (p(correct)), as a function of increasing attentional allocation (neutral, attention+, and attention++; see Methods). Positive Cohen’s *d* indicates inflation (confidence overestimates performance), while negative Cohen’s *d* indicates deflation (confidence underestimates performance). Empirical evidence shows that peripheral inflation is stronger under inattention, so increasing attention ought to reduce inflation strength. Only M4 and M5 correctly predicted that peripheral inflation ought to decrease with increasing attention to the periphery.

|  |  | neutral | attention+ | attention++ |
| --- | --- | --- | --- | --- |
|  |  | 1 | 0.9 | 0.8 |
|  |  | 1 | 0.8 | 0.6 |
| Model | | Cohen's $d$ | | |
| M1 | center | -0.0104 | -0.0236 | 0.0120 |
|  | **periphery** | -0.0054 | 0.0351 | -0.0005 |
| **M2** | **center** | -0.8510 | -1.0903 | -1.5854 |
|  | **periphery** | 1.0273 | 1.0637 | 1.1605 |
| **M3** | **center** | -0.2832 | -0.2572 | -0.1974 |
|  | **periphery** | -0.2212 | -0.2248 | -0.2412 |
| **M4** | **center** | -0.1522 | -0.1474 | -0.1632 |
|  | **periphery** | 0.7287 | 0.6634 | 0.5230 |
| **M5** | **center** | -0.2857 | -0.2536 | -0.2001 |
|  | **periphery** | 0.4846 | 0.4007 | 0.3090 |

In particular, while model M2 (fixed criterion heuristic observer) showed increasingly strong *under*confidence in central vision due to inflation (despite increasing performance), it displayed flat or if anything increasing overconfidence in peripheral vision. If a correction were to be applied such that central Cohen’s *d* for inflation were to remain flat across increasing attentional allocation, this would translate to *increasing* inflation in the visual periphery -- the opposite of empirical reports in the literature. On the other hand, models M4 and M5 showed flat inflation in central vision due to increasing attentional allocation, but if anything slightly *decreasing* overconfidence in the visual periphery. Qualitatively, this behavior matches reports from the literature (Li et al., 2018; Odegaard et al., 2018; Rahnev et al., 2011)(Li et al., 2018; Rahnev et al., 2011).

As a final check of the results of attentional manipulations in comparison to empirical results reported in the literature, we also assessed how attention would change performance and confidence in peripheral locations in a task similar to that used by Odegaard and colleagues (2018) in their Experiment 1. After increasing the possible signal strength to cover a wider range of possible signal and noise combinations to assess the generality of the results (see Methods), we selected from these results only the signal strength and noise combinations in the visual periphery that produced *d’* of about 1.5 in the “attended” condition (attention++; 1.4 < *d’* < 1.6) and about 1.1 (1.0 < *d’* < 1.2) in the “unattended” condition (neutral), consistent with Odegaard et al.’s (2018) findings (see their Figure 2). We then calculated the average confidence in these two conditions separately for correct versus incorrect trials, as done in that paper. We found that only M5 consistently mimicked the results reported by Odegaard and colleagues (2018) by consistently producing no or minimal change in confidence reports for correct trials but a consistent increase in incorrect trials’ confidence (see their Figure 3), both under conditions of the primary simulation ( ) (Figure 4a) and the expanded simulation ( ) (Figure 4b). That is, only M5 showed that under the conditions where *d’* changes from 1.5 to 1.1 as a function of attentional reduction (going from attention++ to neutral), as was found by Odegaard and colleagues (2018), does confidence in correct trials hold exactly constant but confidence in incorrect trials increases.

**Figure 4.**
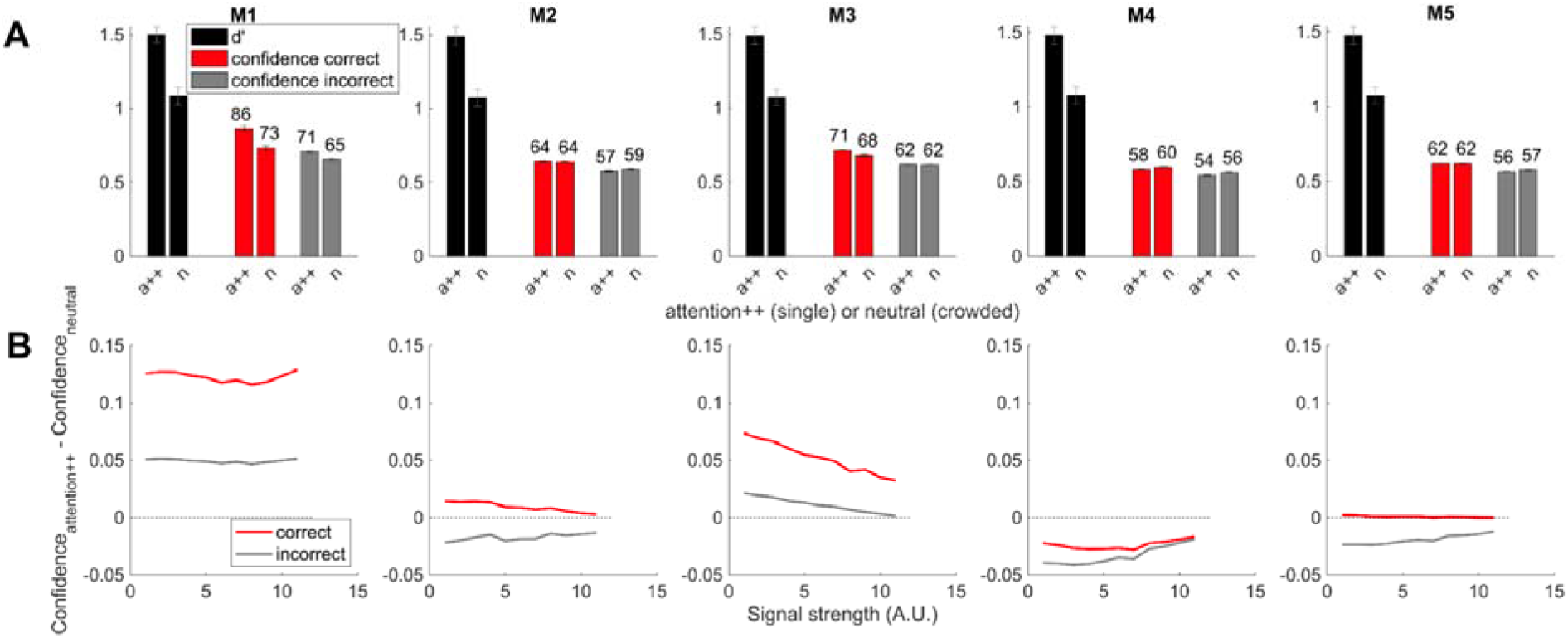
Results of fitting to data reported by Odegaard and colleagues (2018) in their Experiment 1. Those authors reported that reducing attention to a target in the visual periphery (due to a crowding manipulation) led to reductions in performance from d’ ≈ 1.5 to d’ ≈ 1.1, accompanied by no change in confidence for correct trials but an increase in confidence for incorrect trials. We replicated these results by simulating a large range of signal and noise levels, selecting conditions where *d’* was about 1.5 in the attended condition (attention++) and about 1.1 in the unattended condition (neutral), and reading out confidence in the correct and incorrect trials separately. We found that under these *d’* and attention conditions, only M5 produced the target pattern of consistent increases in incorrect confidence but no change in correct confidence both for the conditions of the main simulations (a) and the expanded simulations (b).

## D. DISCUSSION

Here, we asked how knowledge and use of noise statistics across the visual field might inform metacognitive judgments of perceptual decisions. We specifically focused on the phenomenon of visual subjective inflation in the visual periphery -- the phenomenon wherein confidence judgments overestimate decisional accuracy -- including how this phenomenon interacts with attentional manipulations. We hypothesized that the metacognitive system makes use of knowledge of noise statistics in central vision and peripheral vision differently, and that errors in estimation of noise or in use of this prior knowledge might explain why peripheral inflation occurs, and why it becomes stronger under conditions of inattention. Specifically, we proposed that peripheral noise statistics might exhibit a positively-skewed empirical distribution while central noise statistics show a symmetric noise distribution, and that metacognitive heuristics might lead the visual system to systematically overestimate confidence relative to performance in the visual periphery.

We compared five possible model observers which estimate noise and make decisions and confidence in various ways according to Bayesian computations: an observer that possessed no knowledge of natural statistics of noise across the visual field but possessed perfect knowledge of its own noise, and four observers that used such prior knowledge in varying ways. Using Monte Carlo simulations and simple parameter choices as a proof of concept, we showed that Bayesian ideal observer models that either have direct, optimal access to true noise in a sample (model M1: flat Bayesian ideal observer) or optimally estimate noise using accurate empirical noise priors in the center and periphery (i.e., ‘knew’ the peripheral empirical noise distribution was positively skewed; model M3: hierarchical Bayesian ideal observer) predict no over- or underconfidence at all, instead showing confidence behavior that tracks decisional accuracy essentially perfectly. In contrast, heuristic observers that either used a fixed Type 2 criterion for both central and peripheral judgments (model M2: fixed criterion heuristic observer), used the most likely noise level in central or peripheral vision separately (model M4: ‘modes of priors’ heuristic observer), or used a Gaussian estimate for the peripheral noise prior (model M5: hierarchical ‘Gaussian assumption heuristic observer) predicted clear patterns of underconfidence under less noisy conditions and overconfidence under noisier conditions; because of the hypothesized positive skew of empirical noise in the visual periphery, this led these observers to display peripheral inflation. However, only model M2 predicted underconfidence in central vision and overconfidence in the periphery, while M4 and M5 predicted near-perfect confidence in the center and overconfidence in the periphery. Crucially, only M4 and M5 also correctly predicted that increasing sensory and metacognitive precision due to attentional allocation ought to decrease peripheral inflation. And finally, we further expanded these results by conducting qualitative fitting to the empirical findings reported by Odegaard and colleagues (2018); this approach revealed that under the *d’* and attentional manipulation conditions they tested, only M5 produced the consistent increases in confidence in *incorrect* trials but no change in confidence in correct trials, even as *d’* was reduced due to inattention. Together with the general findings for attentional manipulation and peripheral inflation, this pattern of results suggests strongly that M5 may reflect how the system metacognitively estimates noise based on prior experience and incoming information together.

Our results may also explain findings reported in other previous empirical work showing peripheral inflation (Li et al., 2018; Odegaard et al., 2018; Rahnev et al., 2011; Solovey et al., 2014). They also further extend previous signal detection and Bayesian models of peripheral inflation and errors in variance estimation (Fleming & Daw, 2017; Li et al., 2018; Rahnev et al., 2011; Solovey et al., 2014; Zylberberg et al., 2014) to include explicit formulation and exploration of computations that may underlie metacognitive estimates of noise, especially how such mechanisms may rely on natural statistics and previous experience. This formulation thus places the metacognitive system in the same conceptual space as other hierarchical models of perceptual inference which estimate distributions and values for latent variables. Previous work in this area has posited that such hierarchical Bayesian inference might underlie visual inferences about shape by first estimating luminance or specularity (Kersten et al., 2004; Yuille & Bülthoff, 1996), visual inferences about planar slant by first estimating texture isotropy (Knill & Saunders, 2003), multisensory inferences about object heaviness by first estimating relative density (Peters et al., 2016, 2018), or multisensory inferences about numerosity (Shams et al., 2000; Wozny et al., 2008), spatial location (Odegaard et al., 2015; Wozny et al., 2010; Wozny & Shams, 2011), body ownership (Samad et al., 2015), or sensorimotor signal processing (Körding & Tenenbaum, 2007b; Körding & Wolpert, 2003; Wei & Körding, 2011) by first estimating causal relationships among multimodal signals (Körding et al., 2007, 2008; Shams & Beierholm, 2010), among others.

In order to estimate noise in this hierarchical manner, the system would have to have learned the natural statistics of noise as a function of visual eccentricity, and then be able to access and use this expectation in the Bayesian inference process. Such a hypothesis is strongly in line with a body of literature in visual and multisensory perception, suggesting that natural statistics of environmental properties are indeed learned and available to the system in this way. For example, Girschick and colleagues (2011) showed that contour orientations in natural images are more likely to be at cardinal orientations (vertical, horizontal) than oblique orientations (e.g., 45°), and that humans’ visual judgments of orientation are “biased” by these prior expectations in a manner exactly consistent with Bayesian inference in which these expectations serve as the prior. Likewise, humans’ judgments of motion speed are “biased” (again, in a Bayes-optimal way) by the expectation that motion is slow and smooth (Weiss et al., 2002), and Bayes-optimal percepts of heaviness are informed by knowledge of environmental statistics about objects’ densities, with smaller objects more likely to be denser than larger ones (Peters et al., 2015, 2016). These prior expectations about natural statistics also can be learned or modified through statistical learning (Fiser et al., 2010; Gekas et al., 2015; Seriès & Seitz, 2013) in a way that subsequently changes perception (W. J. Adams et al., 2004), and it has been shown that experience with visual scenes can lead to prior expectations of natural statistics being represented in the ongoing spontaneous visual cortical activity of awake, behaving animals (Berkes et al., 2011). That such varied expectations can be learned and used by the perceptual system, and can be updated based on experience, suggests the strong plausibility that the system has learned its own empirical internal noise statistics in an analogous fashion. Our results also suggest, however, that the perceptual system may be using a simplified approximation of these natural statistics, i.e. an assumed symmetric noise prior in the visual periphery. While such simplifications have been demonstrated for other hierarchical priors in the past (e.g., (Peters et al., 2016, 2018)), further work would be necessary to confirm the hypotheses presented here.

The question then becomes, What kind of simplified approximation of natural statistics of noise is the system using? Other previous studies have demonstrated that seemingly suboptimal estimation of noise leading to metacognitive errors may stem from a heuristic estimate of noise rather than a fixed estimate (as in the fixed Type 2 criterion models (Li et al., 2018; Rahnev et al., 2011; Solovey et al., 2014), simple variance misperception (Zylberberg et al., 2014)), or belief errors about noise (Fleming & Daw, 2017). It has also been suggested that a metacognitive update rule which takes into account changing reliability due to sensory noise (Adler & Ma, 2018) or attentional manipulations (Denison et al., 2020) in a simplified linear or quadratic manner rather than fully Bayes-optimal fashion can explain decisional confidence estimates in simple perceptual tasks. These latter studies found that while the fixed criterion type models fail to capture behavior, models that assume the metacognitive system is suboptimally sensitive to changing noise in this particular heuristic fashion can better mimic human behavior. However, these models have not been applied to the differential confidence errors found between central versus peripheral vision, nor do they explain how specifically the shape of the empirical noise priors in central versus peripheral vision (and its mismatch with the observers *beliefs* about such priors) would impact such metacognitive errors. Instead, here we show that Bayesian inference -- especially for model M5, which assumes that the perceptual system possesses an erroneous prior expectation about the shape of the distribution of noise itself -- produces empirically-observed errors in metacognitive estimation as well as the critical pattern of increasing peripheral inflation and selective confidence increases for incorrect trials under increasing inattention. Future work should formally compare the models we explored here with the linear, quadratic, and other heuristic models proposed by other authors in an expanded empirical dataset designed to optimize model fitting and arbitration.

In particular, our results suggest that a candidate cause for peripheral inflation of confidence is a simplified approximation of natural statistics, in the form of an efficient coding scheme for expectations about noise conditioned on visual field location based on empirical priors. In particular, that models M4 and M5 showed peripheral inflation suggests that the system’s prior about noise in the visual periphery may not reflect the true noise experienced, but instead a simplified representation of the most likely noise level to be experienced by the observer in the visual periphery (the mode of the empirical prior) in addition to an impoverished metacognitive ability to estimate such noise in the periphery as compared to the center (*ς*_*center*_ < *ς*_*periphery*_). (Note that an even more extreme simplification, the fixed criterion used by M2, resulted in confidence errors in both center and periphery.) This simplification of prior expectations for complex, skewed, or otherwise non-Gaussian empirical priors into (mixtures of) Gaussians (sometimes formulated as competing priors) has been noted previously (Knill, 2003, 2007; Knill & Saunders, 2003; Yuille & Bülthoff, 1996), and has been suggested to underlie other perceptual illusions (Peters et al., 2016, 2018). Incorrect priors in general, regardless of simplification, can lead to illusions in many areas of perception (Flanagan et al., 2008; Geisler & Kersten, 2002; Teufel et al., 2013; Valton et al., 2019; Weiss et al., 2002), and even if skewed priors are initially learned appropriately it seems the system has difficult using them optimally in cue integration (Acerbi et al., n.d.). Therefore, perhaps due to biological constraints restricting information coding (Heng et al., 2020), we’ve shown here how it is possible that in metacognition, too, efficient, simplified coding of noise expectations in the periphery may lead to overconfidence relative to performance capacity.

Our findings here also suggest fruitful avenues for future study, specifically surrounding the nature and use of the empirical noise prior. First, they suggest that the natural statistics of noise in the visual periphery may be highly positively skewed. Validating this assumption would likely require comprehensive measurement of experienced noise statistics in central versus peripheral vision across a large range of stimulus types. A strong challenge to this endeavor would be that the natural statistic to be measured is *experienced* noise in the visual system rather than natural statistics of noise in the environment. Thus, one promising future approach might be to use established methods for measuring additive and multiplicative internal noise, e.g. through the triple-threshold-versus-contrast function (triple-TVC) approach (Dosher & Lu, 2000, 2017; Lu & Dosher, 1998, 2008) and double-pass procedures (Awwad Shiekh Hasan et al., 2012; Gold et al., 2004; Levi & Klein, 2003; Ratcliff et al., 2018; Vilidaite & Baker, 2017) to quantify internal noise. It would be necessary to conduct these procedures under a range of attentional manipulations, in both the center and visual periphery, and across a large range of tasks *in the same observer* (i.e., a within-subjects design) in order to accurately measure the shape of the internal noise distributions and whether and how they may change conditioned on visual field location or even stimulus type (Bertana et al., 2020). Second, this approach would need to be coupled with established procedures for ‘recovering’ the prior used by an observer in psychophysical tasks in order to compare the true empirical prior to the prior used by the observer. That is, even though such a skewed empirical noise prior may exist in the visual periphery and be recovered by the triple-TVC approach described above, we hypothesized that the system may not rely on the true full prior and instead engage in simplifying heuristic assumptions. While such simplifying heuristics in the face of complex empirical priors have been suggested previously ((Peters et al., 2016, 2018); see also (Yuille & Bülthoff, 1996)), other work suggests even complex, multimodal, non-Gaussian priors can be estimated and used optimally (Girshick et al., 2011; Weiss et al., 2002). Fortunately, a number of methods have been established to ‘recover’ the prior used by an observer in psychophysical estimation tasks. For example, by asking observers to estimate the location of multisensory objects in space, one can engage in model fitting to recover not only a central tendency spatial prior, but also the observer’s prior expectations for whether two stimuli have a common cause (Odegaard et al., 2015, 2017; Wozny et al., 2008) and the decision policy used by the observer (Wozny et al., 2010); likewise, model fitting can recover the prior over motion speed used by observers in a speed estimation task (Weiss et al., 2002). Similar approaches could be used here to recover the noise prior used by observers in central versus peripheral visual estimation tasks, compare findings to the empirical noise priors, and arbitrate among the proposed models more fully. Although this road is daunting, smaller studies may make headway by comparing a few tasks at a time using these paired approaches. Unfortunately, such an endeavor is beyond the scope of the current project, which aims to provide an exploratory proof of concept for how noise priors might be used by the metacognitive system to result in metacognitive errors in noise estimation; we therefore leave such studies for future research.

We admit the present study is limited in its scope due to its nature as proof-of-concept simulations only, without comprehensive parameter fitting to behavioral data; this of course limits its utility, although it provides a principled jumping off point for future empirical studies, as has been done previously (see e.g. (Fleming & Daw, 2017)). These models also do not take into account more nuanced quantitative differences between central versus peripheral vision in terms of other factors, such as crowding or visual search, differential impact by attentional manipulation, and so on (Rosenholtz, 2016; Rosenholtz, Huang, & Ehinger, 2012; Rosenholtz, Huang, Raj, et al., 2012). However, despite the models’ simplicity, the results shown here pave the way for integrating metacognitive noise or uncertainty estimation with a long and established history of hierarchical Bayesian models in perception and cue combination by explicating the specific hypothesis that the metacognitive system builds prior distributions of expected noise that are sensitive not only to experienced ‘environmental’ (within itself) noise statistics but are also sensitive to attentional manipulations, visual field location, and other contextual modulations. We therefore believe that our results provide an important step in realizing the power of such modeling frameworks and empirical approaches for fully explaining how metacognitive computations are performed, and how they may be implemented in neural architecture.

## E. ACKNOWLEDGEMENTS & FUNDING

This work was supported by the Canadian Institute for Advanced Research Azrieli Global Scholars Program in Brain, Mind, & Consciousness (to MAKP). The authors have no other relevant financial or non-financial interests to disclose.

## F. OPEN PRACTICES

Data sharing is not applicable to this article as no datasets were generated or analysed during the current study; the analyses refer to simulations only.

## Notes

### Competing Interest Statement

The authors have declared no competing interest.

### Summary of Updates

This version of the manuscript has been revised to include data fitting of empirical results and to expand and clarify the model descriptions and discussion sections.

